# Ventral pallidum GABA neurons bidirectionally control opioid relapse across rat behavioral models

**DOI:** 10.1101/2022.02.03.479042

**Authors:** Mitchell R. Farrell, Qiying Ye, Yiyan Xie, Jeanine Sandra D. Esteban, Stephen V. Mahler

## Abstract

Opioid addiction is a chronic, relapsing disorder. Whether addicted individuals are forced to abstain or they decide themselves to quit using drugs, relapse rates are high—especially upon encountering contexts and stimuli associated with prior opioid use. Rodents similarly show context- and cue-induced reinstatement of drug seeking following abstinence, and intriguingly, the neural circuits underlying these relapse-like behaviors differ when abstinence is involuntarily imposed, versus when animals decide themselves to cease taking drug. Here, we employ two complementary rat behavioral models of relapse-like behavior for the highly reinforcing opioid drug remifentanil, and asked whether GABAergic neurons in the ventral pallidum (VP^GABA^) control opioid seeking under these behavioral conditions. Specifically, we asked how chemogenetically stimulating VP^GABA^ neurons with clozapine-N-oxide (CNO) influences the ability of contextual or discrete remifentanil-paired cues to reinstate drug seeking following either voluntary (punishment-induced; Group^Punish^), or experimenter-imposed (extinction training; Group^Ext^) abstinence. In Group^Punish^ rats, we also chemogenetically inhibited VP^GABA^ neurons, and examined spontaneous VP activity (Fos) during cued-reinstatement. In both Group^Punish^ and Group^Ext^ rats, stimulating Gq-signaling in VP^GABA^ neurons augmented remifentanil reinstatement in a cue- and context-dependent manner. Conversely, engaging inhibitory Gi-signaling in VP^GABA^ neurons in Group^Punish^ suppressed cue-induced reinstatement, and additionally cue-triggered seeking was correlated with Fos in rostral, but not caudal VP. In contrast, neither stimulating nor inhibiting VP^GABA^ neurons influenced unpunished remifentanil self-administration. We conclude that VP^GABA^ neurons bidirectionally control opioid seeking regardless of the specific relapse model employed, highlighting their fundamental role in opioid relapse-like behavior across behavioral models, and potentially across species.

**Highlights:**

- We acutely inhibit or stimulate VP GABA neurons during opioid seeking
- VP GABA neurons mediate relapse-like behavior across behavioral models
- Behavioral context impacts DREADD stimulation of behavior, not VP activity
- Rostral, not caudal VP Fos correlates with opioid reinstatement

## Introduction

Opioid addiction is a disorder characterized by persistent drug use despite adverse consequences, and chronic risk of relapse after quitting. Though addicted individuals frequently quit drug use due to mounting negative consequences, they often still relapse despite their desire to remain abstinent [1–3]. In particular, exposure to drug-associated cues or contexts often elicit cravings and promote relapse [4].

Much preclinical work has examined the neural circuits of cue-, stress-, or drug-induced relapse-like behavior in rodents, especially using models involving experimenter-imposed abstinence prior to reinstatement of drug seeking [5]. Yet these conventional models do not capture the voluntary initiation of abstinence that is typical of addicted humans seeking to control their use in the face of mounting negative life consequences—in fact, rats in these experiments have little disincentive to pursue drugs when it is available. The presence of such disincentives to drug use might be important for preclinically modeling addiction-related behaviors, especially since the neural substrates underlying cue-induced reinstatement differ when rats previously chose to stop taking drugs, rather than being forced to do so with extinction training [6,7]. Clearly, no single rodent model captures all aspects of the human use-cessation-relapse cycle, so to maximize likelihood of translational relevance, we propose that putative addiction interventions should be tested in multiple rodent behavioral models including those optimizing human relevance [8]. We hope that by understanding the neural circuits underlying relapse-like behaviors across animal models that capture distinct features of the human disorder, we can identify more promising candidates for targeting brain-based psychiatric intervention in heterogeneous humans struggling to control their drug use.

Many prior rodent reinstatement studies have examined the brain substrates of opioid relapse following experimenter-imposed homecage abstinence (such as incubation of craving), or extinction training [9–16]. Fewer studies have used models in which rodents instead voluntarily cease their drug use, for example due to delivery of punishing shocks co-administered with drug. For opioid drugs, this is partly due to the methodological consideration that the analgesic properties of opioid drugs can diminish the ability of shock to suppress drug seeking. Here, we circumvented this problem by using the short-acting, but highly reinforcing μ opioid receptor agonist remifentanil, similar to a model presented by Panlilio and colleagues [6,17]. Since remifentanil is rapidly metabolized [18], we were able to develop a shock-based voluntary abstinence/reinstatement procedure, allowing for direct comparison of opioid reinstatement following either voluntary (punishment-based), or imposed (extinction-based) abstinence.

Specifically, we examined the role of ventral pallidum (VP), a brain region tightly embedded within mesocorticolimbic motivational circuits where opioid signaling plays important roles in reward-related processes [19,20]. Locally applied μ opioid receptor agonists in VP induce robust food intake and locomotion, and enhance pleasure-like reactions to sweet tastes [20,21]. Systemically administered heroin or morphine decrease extracellular GABA levels in VP [22,23], indicating the region is also involved in effects of systemically-administered addictive opioid drugs. Accordingly, lesioning or inactivating VP neurons diminishes high-effort responding for heroin [24] and the ability of heroin priming injections reinstate heroin seeking following extinction training [25]. VP is also required for high-effort intake of remifentanil, since local application of an orexin receptor antagonist attenuates remifentanil motivation in both behavioral economic and cue-induced reinstatement tasks [26]. Clearly, VP is a key node in the circuits underlying the rewarding and relapse-inducing effects of addictive opioid drugs.

This said, VP is a heterogeneous structure, and little is known about how this functional heterogeneity impacts relapse-like behavior. VP contains subpopulations of neurons with different neurotransmitter profiles and behavioral functions [27–32], and rostrocaudal as well as mediolateral functional heterogeneity are also apparent [33–42]. For example, the rostral portion of VP is critical for cue-induced cocaine seeking, whereas its caudal aspect is instead required for cocaine-primed reinstatement [33]. Caudal VP also contains a ‘hedonic hotspot’ wherein local application of a selective μ opioid receptor agonist (or orexinA peptide [43]) selectively enhances taste pleasure [34,44].

VP GABAergic (VP^GABA^) neurons, which span both rostral and caudal VP zones, appear to play a specialized role in reward-related processes, in contrast to intermingled VP glutamate neurons, which instead mediate aversive salience processes [27,28,30,31,45]. For example, mice find optogenetic stimulation of VP^GABA^ neurons reinforcing, and these neurons show endogenous firing patterns consistent with the encoding of incentive value of rewards and reward-predictive cues [27,30,46]. Stimulation of a subset of VP^GABA^ neurons expressing enkephalin also increases reinstatement of cocaine seeking in mice following experimenter-imposed extinction training [29], and inhibiting VP^GABA^ neurons suppresses context-induced alcohol seeking after extinction training in rats [47]. Though these findings point to an important role for VP^GABA^ neurons in highly-motivated and relapse-relevant behaviors, no studies have yet examined the role of these neurons in opioid seeking, nor compared their functions in relapse models capturing dissociable addiction-relevant behavioral processes.

Here we address this gap by determining how VP^GABA^ neurons regulate remifentanil intake and seeking using two distinct models of relapse-like behavior, including a newly developed voluntary abstinence-based reinstatement task. Using DREADDs [48], we found that inhibiting VP^GABA^ neurons decreased opioid relapse after voluntary abstinence, whereas stimulating VP^GABA^ neurons strongly increased opioid seeking regardless of the way in which abstinence was achieved prior to reinstatement. Moreover, chemogenetic effects largely relied on the presence of response-contingent cues, suggesting that VP^GABA^ neurons may play a special role in discrete cue-induced opioid seeking. Consistent with these findings, the degree of endogenous VP activity correlated with opioid reinstatement behavior in individual animals, but only in its rostral, but not caudal, subregion. Further, we found that neither inhibiting nor stimulating VP^GABA^ neurons influenced unpunished remifentanil self-administration, highlighting a selective role for these neurons in relapse-like drug seeking, rather than in the primary reinforcing effects of remifentanil. Together, these results point to a fundamental and specific role for VP^GABA^ neurons in opioid drug relapse-like behavior in rats, regardless of the behavioral model employed. These results beg the question of whether VP is similarly involved in human drug relapse, and if so, whether such circuits might be a promising future target for clinical treatment of opioid addiction.

## Materials and Methods

### Subjects

GAD1:Cre transgenic rats (*n* = 32 males, *n* = 9 females) and wildtype littermates (*n* = 22 male, *n* = 12 female) were used as subjects. They were pair-housed in temperature, humidity, and pathogen-controlled cages under a 12:12 hr reverse light/dark cycle, and were provided ad libitum food and water in the homecage throughout all experiments. Experiments were approved by University of California Irvine’s Institutional Animal Care and Use Committee, and were conducted in accordance with the NIH Guide for the Care and Use of Laboratory Animals [49].

### Surgery

Procedures for GAD1:Cre-dependent DREADD viral injections in VP were conducted as previously described [50]. Briefly, anesthetized GAD1:Cre rats and wildtype littermates were injected with one of three AAV2 viral constructs obtained from Addgene: hSyn-DIO-hM4D(Gi)-mCherry, hSyn-DIO-hM3D(Gq)-mCherry, or hSyn-DIO-mCherry (~0.3 μL/hemisphere, titers: 1.2 x 1013 GC/mL). During the same surgery, rats were implanted with indwelling, back-mounted right jugular vein catheters for chronic drug self-administration as previously described [33,51–53].

### Drugs

Frozen powder aliquots of clozapine-N-oxide (CNO; NIDA) were diluted in 5% dimethyl sulfoxide (DMSO; Sigma-Aldrich), vortexed for 10 s, then diluted with sterile 0.9% saline to a concentration of 5 mg/mL. CNO was mixed fresh on each test day, and injected at 5 mg/kg (i.p.) in all experiments, 30 min prior to the start of behavioral testing. Vehicle solutions were 5% DMSO in saline, injected at 1 mL/kg. Rats were surgically anesthetized with ketamine (56.5 mg/kg) and xylazine (8.7 mg/kg), and given the non-opioid analgesic meloxicam (1 mg/kg). Remifentanil hydrochloride was dissolved in sterile 0.9% saline to a concentration of 38 μg/mL for self-administration.

### Group^Punish^ training

#### Self-administration phase

Following recovery from surgery, hM4Di (*n* = 8 males, 5 females), hM3Dq (*n* = 13 male, *n* = 0 female) and control rats (*n* = 9 males, 8 females) were initially trained in a distinct Context A (peppermint odor, white light, and bare walls) during 2 hr daily sessions. They learned to press an active lever for intravenous infusions of remifentanil (1.9 μg/50 μL/ infusion), a short-acting μ opioid receptor agonist [18,54], accompanied by a light + tone cue (3.6 s stimulus light + 2.9 kHz tone). Infusions/cues were followed by a 20 s timeout period, signaled by dimming of the houselight, during which lever presses were unreinforced, but recorded. Presses of an inactive lever positioned on the opposite side of the chamber were without consequence. Training in Context A proceeded on the following schedules of reinforcement: 5-6 days of fixed-ratio 1 (FR1), 2 days of variable interval 5 (VI5), 2 days of VI15, and finally 5 days of VI30.

#### Punishment training

Next, Group^Punish^ rats were moved to a distinct Context B (orange odor, red light, and polka dot walls), where active lever presses (on a VI30 schedule) yielded the same dose of remifentanil and cues. However, in Context B infusions were accompanied by a 50% probability of footshock, delivered concurrently with the start of the infusion/cue. All rats were initially given one drug-free punishment training day in Context B (0.30 mA footshock intensity), in order to determine the degree of punishment-induced suppression of self-administration in each individual. An initial cohort (*n* = 7 hM4Di, *n* = 5 hM3Dq, *n* = 4 controls) was then used to examine effects of inhibiting or stimulating VP^GABA^ neurons during punished remifentanil self-administration. This group was administered CNO (5 mg/kg) or vehicle on each subsequent daily punishment training day according to the following protocol: 2 days with 0.30 mA shocks, followed by 2 days each at footshock intensities increasing by 0.15 mA on each step, up to a maximum of 1.65 mA, to suppress pressing. Punishment training ceased in all rats upon reaching voluntary abstinence criterion (< 25 AL presses on 2 consecutive days, days to criterion mean ± SEM: 16.8 ± 0.53). 48+ hours after abstinence criterion was reached in these rats, they were given a final Context B punished self-administration test without veh/CNO, to measure maintenance of abstinence in the absence of VP manipulation.

Since no signs of CNO effects were observed on punished drug seeking in this cohort (data not shown), subsequent cohorts of rats (*n* = 27) received a modified protocol aimed at more rapidly inducing voluntary abstinence, without daily CNO/vehicle treatment. These rats were trained on the Context B punished self-administration procedure according to the following protocol: 1 day of 0.30 mA shocks, followed by 1 day each at 0.45, 0.60, 0.75, 0.90, and 1.05 mA. Rats trained with both protocols reached the same voluntary abstinence criterion (< 25 AL presses on 2 consecutive days), and had similar levels of pressing by the end of training (average active lever presses on last 2 days in the 2 cohorts: *t*_41_ = 0.15, *p* = 0.88). Both cohorts also showed similar levels of subsequent reinstatement behavior (two-way ANOVA on reinstatement type x cohort; no main effect of cohort: F_(1, 121)_ = 0.28, *p* = 0.60; or reinstatement type x cohort interaction: F_(2, 121)_ = 1.68, *p* = 0.19). Therefore groups were collapsed for subsequent analyses of DREADD effects reinstatement in Group^Punish^.

#### Reinstatement testing

After achieving abstinence criterion in Context B, all Group^Punish^ hM4Di, hM3Dq, and control rats were then administered a series of reinstatement tests to determine how inhibiting or stimulating VP^GABA^ neurons affected reinstatement in Contexts A and B, with or without response-contingent cues (and without further remifentanil). Counterbalanced vehicle and CNO injections were administered in a within subjects design prior to each reinstatement conditions, in the following order, with 48 hrs between each test: 1) Context B with response-contingent cues (*n* = 42) and 2) with no cues (*n* = 42) and 3) Context A with (*n* = 43) and 4) with no cues (*n* = 27). Note that a subset of rats (*n* = 16) did not undergo the Context A with no cues tests, due to a Spring of 2020 COVID-19 shutdown. All tests occurred in the absence of remifentanil, and active/inactive lever presses were recorded.

#### Remifentanil self-administration testing

Following reinstatement testing, a subset of Group^Punish^ rats (*n* = 5 male, *n* = 1 female hM4Di, *n* = 8 male, *n* = 0 female hM3Dq, *n* = 6 male, *n* = 6 female controls) were retrained to self-administer remifentanil and light + tone cue in a distinct chamber (ie, neither Context A nor B) on a VI30 schedule, identical to initial training. Counterbalanced vehicle and CNO tests were administered upon achieving stability criterion (<25% change in active lever presses on 2 consecutive days), with at least one day of restabilization between tests.

### Group^Ext^ training

A separate cohort of hM3Dq rats (*n* = 8 males, 4 females) and controls (*n* = 11 males, 4 females) were trained to self-administer remifentanil exactly as was Group^Punish^: 14 daily 2 hr sessions of up to VI30 remifentanil/cue self-administration occurred in Context A. Next, Group^Ext^ also moved to Context B, but for this group active lever presses (VI30 schedule) delivered no drug infusions, cues, or shocks (extinction conditions), unlike in Group^Punish^ where Context B active lever presses yielded all three. Extinction training continued in Context B for Group^Ext^ rats until the criterion was met (< 25 active lever presses on 2 consecutive sessions). After lever pressing was extinguished, Group^Ext^ rats then underwent CNO/vehicle tests on each of the 4 reinstatement types, as described for Group^Punish^ rats above: 1) Context B with cues (*n* = 27) and 2) with no cues (*n* = 27) and 3) Context A with cues (*n* = 27) and 4) with no cues (*n* = 27).

### hM3Dq-DREADD Fos validation

Our prior work validated the function of hM4Di-DREADDs in VP^GABA^ neurons of GAD1:Cre rats [50]. Here, we confirmed the function of hM3Dq-DREADDs in this model, using Fos as a marker of neural activity. To do so, two experimentally-naïve groups were first tested. The first group expressed mCherry in VP^GABA^ neurons (mCherry-only, *n* = 3), and the second group instead expressed hM3Dq-mCherry in VP^GABA^ neurons (*n* = 3). Both groups were injected with CNO before returning to the homecage for 2.5 hrs, then were perfused for analysis of Fos in mCherry-expressing VP^GABA^ neurons.

Further, we also asked whether hM3Dq-induced Fos differed depending on the behavioral situation the rat was in. In a final 2 hr session, we therefore stimulated VP^GABA^ neurons of hM3Dq-mCherry Group^Ext^ rats previously tested for reinstatement (described above). These rats were injected with CNO, then 30 min later we noncontingently presented 66 evenly spaced remifentanil-paired cues (*n* = 3), or no cues (*n* = 4) over 2 hrs in a novel operant chamber (ie, neither Context A nor B), without levers extended. This number of cues was selected as it was the average number of cues delivered by rats during self-administration training. Rats were perfused immediately after this final cue/no-cue session for analysis of Fos in mCherry-expressing VP^GABA^ neurons.

### Immunofluorescent and immunohistochemical staining

#### Immunofluorescent visualization of DREADD expression

To visualize DREADD localization in each subject, VP sections were stained for substance P, which delineated VP borders from surrounding basal forebrain [32,55], and mCherry, which labeled DREADD-expressing GABA neurons. Rats were perfused with 0.9% saline and 4% paraformaldehyde, brains were postfixed for 16 hrs, then cryoprotected in 20% sucrose-azide. Brains were sectioned at 40 μm using a cryostat, and 6-8 sections spanning VP’s rostrocaudal axis (from bregma +0.7 to bregma −0.6) were collected and stained, as described previously (Farrell et al., 2021). Briefly, sections were first blocked in 3% normal donkey serum (NDS), then incubated overnight in rabbit anti-substance P (ImmunoStar; 1:5000) and mouse anti-mCherry antibodies (Clontech; 1:2000) in PBST-azide with 3% NDS. Finally, sections were incubated for 4 hrs in Alexafluor donkey anti-Rabbit 488 and donkey anti-Mouse 594 (Thermofisher). Sections were mounted, coverslipped with Fluoromount (Thermofisher), and imaged at 5x magnification with a Leica DM4000 with StereoInvestigator software (Microbrightfield). Viral expression sites were mapped in each rat referencing a rat brain atlas [56] and observed VP borders.

#### Endogenous reinstatement-related VP Fos visualization

A subset of Group^Punish^ rats were perfused directly following a final 2 hr reinstatement test (Context A with cues: *n* = 9, Context B with cues: *n* = 8), in the absence of CNO/Veh injection. To quantify VP neural activity following this test, we stained a set of slices throughout VP for Fos protein as a marker of neural activity, and co-stained the same samples for substance P to define VP borders on each section. Tissue was blocked in 3% NDS, incubated overnight in rabbit anti-Fos primary antibody (Millipore, 1:10000), then for 2 hrs in biotinylated donkey anti-rabbit secondary antibody (Jackson Immuno, 1:500), followed by 90 min amplification in avidin-biotin complex (ABC; Vector Lab, 1:500). Sections were then reacted in 3,3’-Diaminobenzidine (DAB) with nickel ammonium sulfate, to reveal a black nuclear stain for Fos protein. After washing, sections were incubated overnight in mouse anti-substance P (Abcam, 1:10000), then donkey anti-mouse biotinylated secondary antibodies (Jackson Immuno, 1:500), then amplified with ABC. Another DAB reaction without nickel was conducted, yielding a light brown product visualizing substance P-immunoreactive processes and neuropil.

#### mCherry and Fos visualization for hM3Dq-DREADD validation

Sections were stained using double DAB immunohistochemistry to visualize neurons expressing Fos (black nuclei) and mCherry (brown soma). Procedures mirrored above, except after the Fos stain, a mouse anti-mCherry primary antibody (Takara Bio, 1:5000) was used instead of the substance P primary antibody to visualize hM3Dq-mCherry or mCherry-expressing cells.

### Fos quantification

#### Substance P and Fos co-stain quantification

To examine reinstatement-related Fos within VP borders, stained sections were mounted, coverslipped, and imaged at 10x magnification, and using ImageJ two observers blind to experimental conditions manually counted all Fos+ nuclei within the substance-P defined VP borders on 4 sections/rat. These sections spanned the rostrocaudal extent of VP, from bregma +0.7 to bregma −0.6. Fos counts from the left and right hemisphere were averaged for each section, and these section averages were averaged to generate a per rat mean Fos value, which was used for statistical analyses. In addition, sections were divided into rostral and caudal bins (1-3 sections/bin) in accordance with their location relative to bregma (rostral VP >0 AP relative to bregma, caudal VP ≤ 0 AP relative to bregma). An inter-rater reliability measure showed a strong positive correlation between the two observers’ per rat average Fos quantification (Pearson’s correlation: *r* = 0.94, *p* < 0.0001).

#### mCherry and Fos co-stain quantification

To examine hM3Dq-induced Fos in VP^GABA^ cells, stained sections were mounted, coverslipped, and imaged at 10x magnification. mCherry+ cells, Fos+ cells, and mCherry+/Fos+ cells within VP borders (estimated based on [56]) were counted in ImageJ. Two sections per rat were quantified from near the center of VP injection sites, and counts from the left and right hemisphere of each section were averaged. The two section averages were then combined to generate a per rat mean, which was used for statistical analyses of group effects.

### Data analysis

Data were analyzed in Graphpad Prism. Repeated measures two-way ANOVAs with lever (inactive, active) and treatment (vehicle, CNO) were conducted for each set of reinstatement and self-administration tests, accompanied by Sidak post hoc tests. One-way or two-way ANOVAs were used to examine differences in lever pressing among vehicle-treated rats during their reinstatement tests, coupled with Sidak post hoc tests. Repeated measures two-way ANOVAs with DREADD type (Gi, Gq, control) and treatment (vehicle, CNO) were used to compare active lever pressing for each reinstatement condition to confirm DREADD-specificity of CNO effects. A one-way ANOVA was used to compare Fos across reinstatement conditions, coupled with Sidak post hoc tests. Paired t-tests were used to compare final-day self-administration behavior to first day punishment behavior. Pearson’s correlations were used to examine the relationship between VP Fos and active lever pressing on a final reinstatement test, as well as inter-rater reliability between blinded observers. Log-rank (Mantel-Cox) test compared the number of days required to reach abstinence criterion for Group^Punish^ versus Group^Ext^. One rat in Group^Ext^, and 1 in the substance P/Fos reinstatement experiment were removed from analyses as outliers (>3 standard deviations from the mean). Statistical significance thresholds for all analyses were set at *p* < 0.05, two-tailed.

## Theory

Opioid addiction is a chronically-relapsing disorder, but we have few effective therapies to offer those trying to remain abstinent from opioid drugs. In part, this may be due to 1) a lack of understanding of the neural circuits underlying compulsive use and relapse to opioid seeking, and 2) a failure to capture key aspects of the addiction/relapse cycle in preclinical addiction models. Here, we examine the activity, necessity, and sufficiency of VP^GABA^ neurons for reinstatement of opioid drug seeking using complementary rat behavioral models of relapse, and provide converging evidence for them playing a critical role in opioid relapse.

## Results

### Training in Context A and response suppression in Context B via punishment or extinction training

During Context A training, Group^Punish^ and Group^Ext^ exhibited comparable levels of remifentanil self-administration (Figure S1A; infusions obtained throughout training: *t*_68_ = 1.30, *p* = 0.20). In Group^Punish^, as we previously saw using an analogous cocaine model [51], shifting from unpunished Context A to Context B where 50% of infusions were met with contingent footshock decreased active lever responding (last day Context A vs. 1^st^ day Context B: active lever, *t*_42_ = 2.61, *p* = 0.022), and increased inactive lever pressing (*t*_42_ = 2.70, *p* = 0.0098). In Group^Ext^, shifting from Context A to Context B also decreased active lever responding (last day self-administration vs. 1^st^ day extinction active lever: *t*_26_ = 3.16, *p* = 0.004) and increased inactive lever responding (*t*_26_ = 5.22, *p* < 0.0001). Across Context B training, Group^Punish^ rats suppressed their active lever pressing to criterion in fewer days than Group^Ext^ rats (Fig S1B, Log-rank Mantel-Cox survival analysis test, χ^2^ = 18.52, *p* < 0.0001).

### Cues and contexts gate remifentanil reinstatement in both Group^Punish^ and Group^Ext^ rats

In Group^Punish^ rats, opioid reinstatement was impacted by both context and the presence or absence of discrete, response-contingent cues (Fig 1B, one-way ANOVA on vehicle test day active lever pressing: F_(3, 150)_ = 10.66, *p* < 0.0001). In Context A, more seeking was seen with cues than without (Sidak post hoc: *p* = 0.0005), and cues elicited more pressing in Context A than they did in Context B (*p* = 0.0006). Context A with cue reinstatement was also greater than in Context B without cues (*p* < 0.0001). Pressing was similar in Context A without cues to pressing in Context B, with or without cues (*p*s > 0.83).

**Figure 1.**
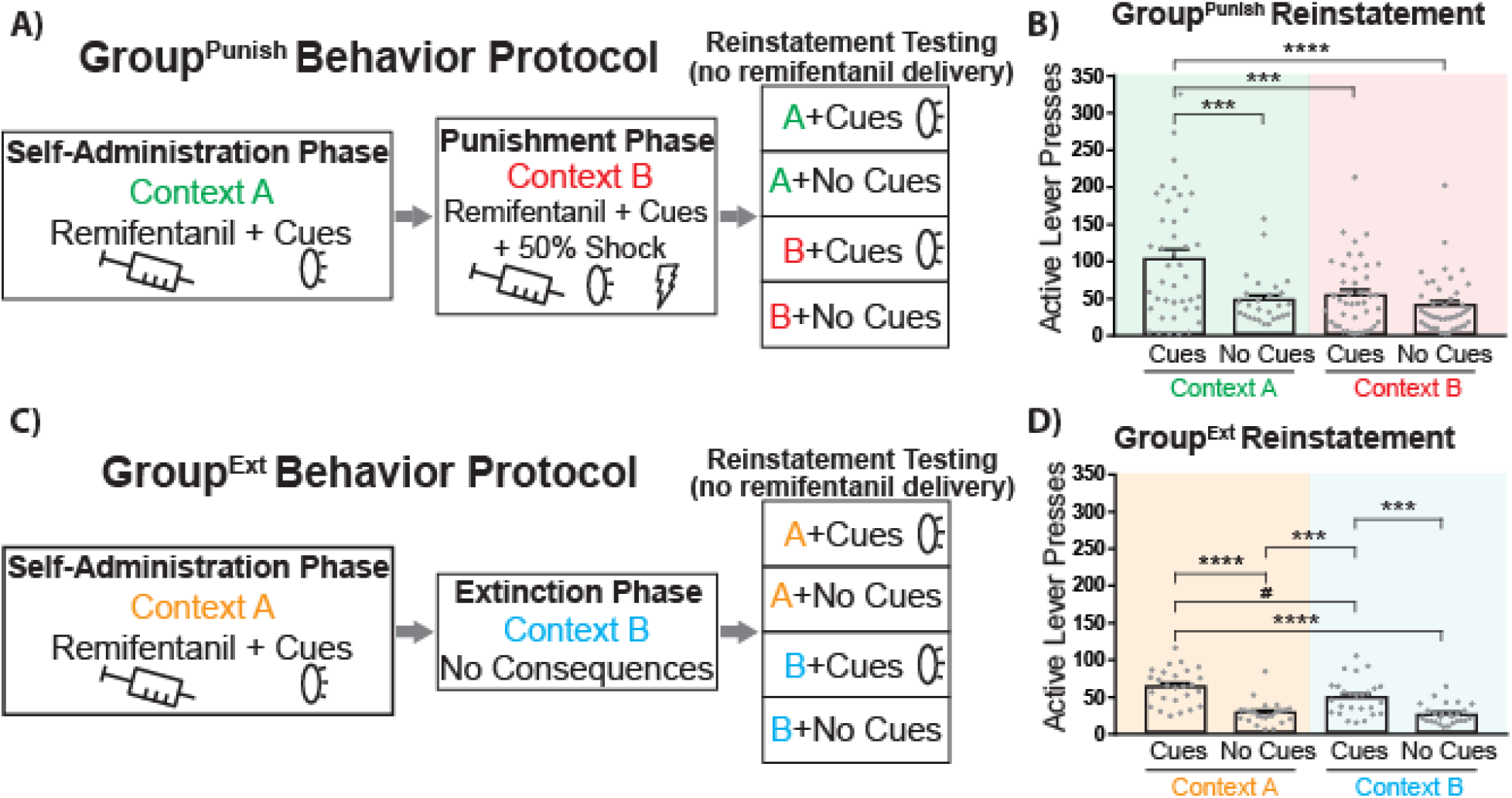
Behavioral testing schematic, and vehicle-day reinstatement following punishment-versus extinction-induced abstinence. **A)** Schematic of the behavioral training for Group^Punish^ rats undergoing self-administration in Context A, punishment in Context B, and a series of reinstatement tests in both contexts, with or without response-contingent cues. **B)** In Group^Punish^ rats, active lever pressing in Context A (green shading) and punishment Context B (red shading), with or without response-contingent discrete cues is shown for vehicle test days. **C)** Schematic of the behavioral training for Group^Ext^ rats undergoing self-administration in Context A, extinction in Context B, and a series of reinstatement tests in both contexts, with or without response-contingent cues. **D)** In Group^Ext^ rats, active lever pressing in Context A (orange shading) or extinction Context B (blue shading), with or without discrete cues is shown for vehicle test days. Individual rats shown as gray dots. One-way ANOVA, Sidak post hoc: *p*# < 0.06, *p*\*\* < 0.01, *p*\*\*\* < 0.001, *p*\*\*\*\* ≤ 0.0001. Data presented as mean ± SEM.

In Group^Ext^ rats, opioid reinstatement was also impacted by both context and discrete, response-contingent cues (Fig 1D, repeated measures ANOVA: F_(3, 75)_ = 30.55, *p* < 0.0001). Pressing was greater in the Context A with cues test than in either context without cues (Sidak post hocs; Context A with cues versus no-cue tests in Context A: *p* < 0.0001; or Context B: *p* < 0.0001), and cue-elicited pressing trended toward being greater in Context A than in Context B (*p* = 0.058). In Context B, pressing was greater with cues than without them (*p* = 0.0001).

Overall, reinstatement in Group^Punish^ was greater than reinstatement in Group^Ext^ [vehicle day data; two-way ANOVA with group (Group^Punish^, Group^Ext^) and reinstatement condition (Context A with/without cues, Context B with/without cues) as factors: F_(1, 250)_ = 11.44, *p* = 0.0008]. This effect was in part due to high levels of pressing of Group^Punish^ rats in Context A with cues, as there was greater reinstatement in Context A with cues relative to all other reinstatement conditions in both Group^Punish^ and Group^Ext^ (Sidak post hocs: ps < 0.016). These results indicate that response-contingent cues reinstate seeking following either punishment or extinction training, but the modulation of this by context may be greater in Group^Punish^, relative to Group^Ext^.

### Rostral VP neural activity is positively correlated with cue-induced reinstatement

To determine whether VP Fos was associated with reinstatement behavior, a subset of Group^Punish^ rats, following all 8 reinstatement tests with vehicle and CNO, were sacrificed following a final reinstatement test in Context A with cues, Context B with cues, or directly from their homecage. Greater Fos expression was found in both Context A (mean ± SEM: 250.97 ± 27.71) and Context B-tested rats (m = 315.77 ± 47.87), relative to homecage-tested controls (m = 105.06 ± 18.15; one-way ANOVA, F_(2, 21)_ = 12.25, *p* = 0.0003; Sidak post hoc: safe with cues vs. homecage, *p* = 0.0079; punishment with cues vs. homecage, *p* = 0.0003). No difference in VP Fos expression was detected between Context A with cues and Context B with cues (Sidak post hoc: *p* = 0.43). Fos in rostral (Fig 2A), but not caudal (Fig 2B), VP correlated with total active lever presses during the final reinstatement test (*r* = 0.62, *p* = 0.014), consistent with our prior report showing that rostral VP Fos is associated with cue-induced cocaine seeking [33].

**Figure 2.**
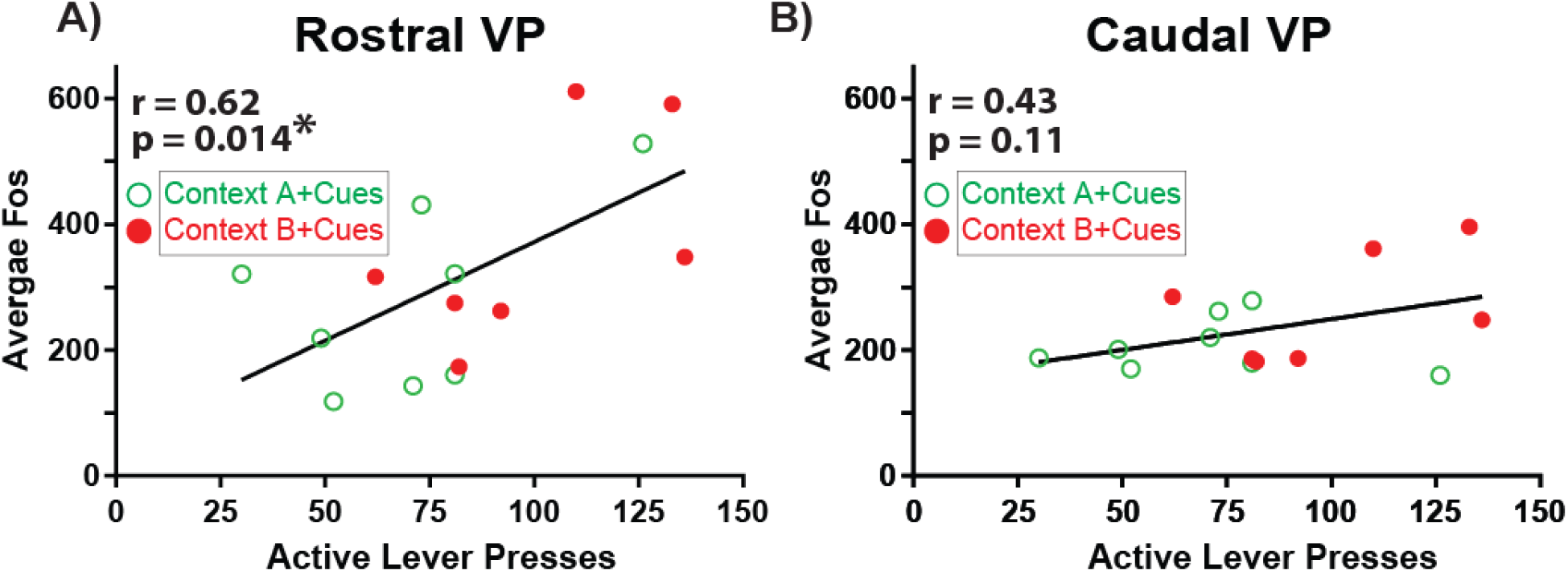
Rostral, but not caudal, VP Fos correlates with remifentanil seeking. **A)** In Group^Punish^ rats, Fos in rostral VP (anterior of bregma) positively correlates with cue-induced opioid reinstatement (active lever presses). Green circles represent rats tested with cues in Context A, red dots represent those tested with cues in Context B. **B)** Fos in caudal VP (posterior of bregma) was uncorrelated with opioid reinstatement. Pearson correlation: p* < 0.05.

### hM4Di- and hM3Dq-DREADD expression in VP^GABA^ neurons

GAD1:Cre rats expressing DREADDs in >50% of VP (defined by substance P) were included for analyses, for a total of 13 GAD1:Cre hM4Di-expressing (hM4Di) rats (Fig 3A-B, Group^Punish^: *n* = 8 males, 5 females) and 25 GAD1:Cre hM3Dq-expressing (hM3Dq) rats (Fig 3C-D, Group^Punish^; *n* = 13 males, 0 females; Group^Ext^: *n* = 8 males, 4 females). Control rats were designated as those with DREADD expression outside of VP (*n* = 2), mCherry expression (*n* = 8), or Cre- rats with no expression (*n* = 10).

**Figure 3.**
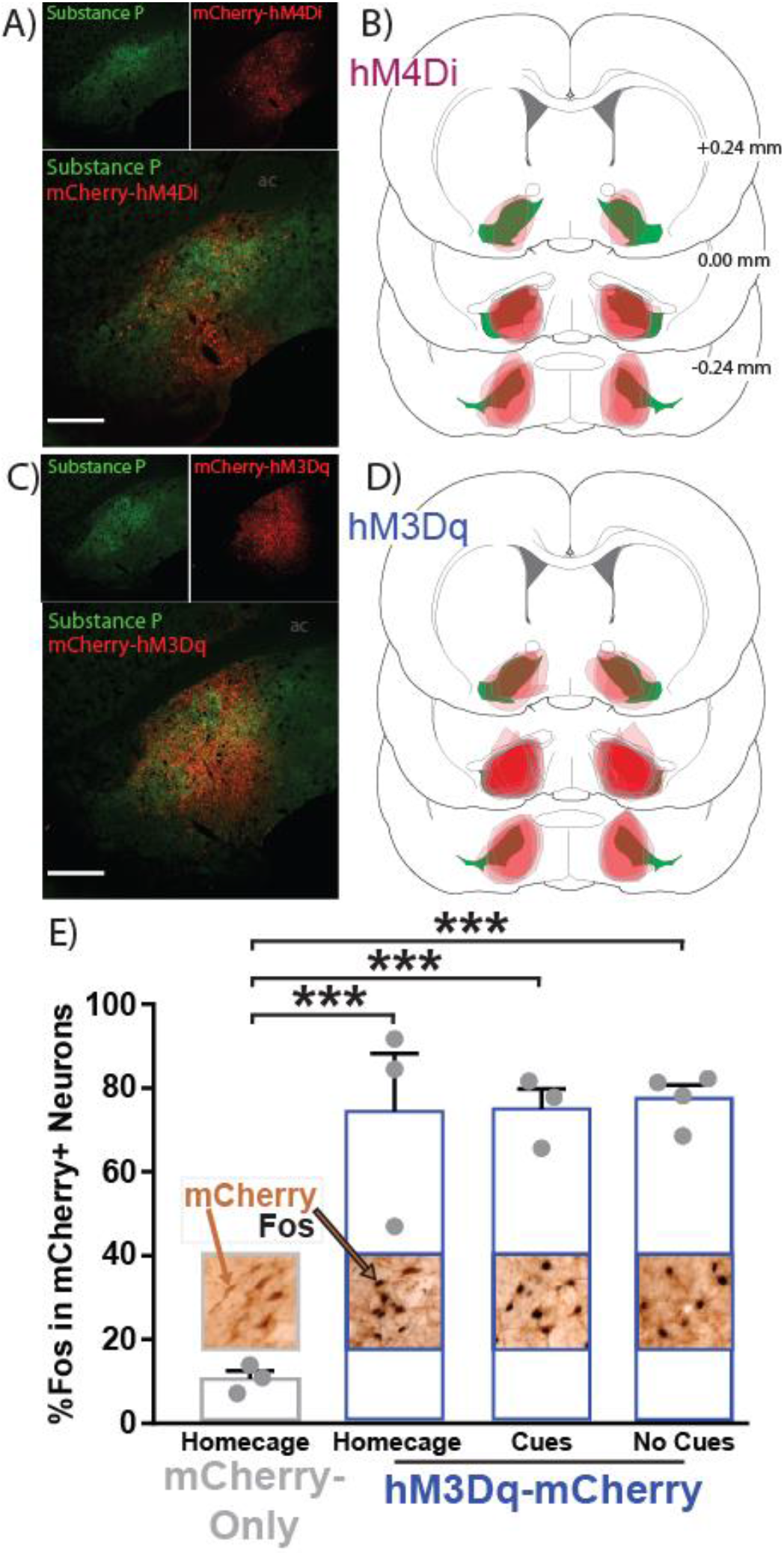
VP^GABA^ DREADD localization and hM3Dq validation. **A)** Expression of hM4Di-mCherry (red) localized largely within VP borders defined by substance P (green). **B)** Coronal sections depicting the center of hM4Di-mCherry expression (red) for each rat along VP’s rostrocaudal axis relative to bregma (substance P-defined VP borders = green). **C)** Expression of hM3Dq-mCherry (red) is similarly localized within VP borders (green). **D)** Coronal sections similarly depicting the center of hM3Dq-mCherry expression (red) for each rat is shown. **E)** CNO treatment in hM3Dq-mCherry rats tested in the homecage exhibited greater Fos in mCherry+ neurons (2^nd^ bar, blue), than in homecage mCherry-only rats treated with CNO (1^st^ bar, gray). CNO-treated hM3Dq-mCherry rats exposed to remifentanil-paired cues (3^rd^ bar, blue) or no cues (4^th^ bar, blue) had more Fos+ mCherry neurons homecage rats (1^st^ bar, gray). However, the hM3Dq stimulation of Fos was no different in the presence or absence of cues. Images above/embedded within bars depict 10x images of immunohistochemical staining of mCherry (brown) within VP borders, and Fos+ nuclei (black). Example mCherry-only and mCherry+Fos double-labeled neuron indicated with brown and brown/black arrows, respectively. Individual rat data shown as gray dots on top of bars. One-way ANOVA, Sidak post hoc: *p*\*\*\* ≤ 0.001. Data presented as mean ± SEM.

### CNO increases Fos immunoreactivity in hM3Dq neurons

CNO treatment induced more Fos in hM3Dq-expressing neurons, relative to mCherry-only neurons (Fig 3E, One-way ANOVA: F_(3, 9)_ = 20.56, *p* = 0.0002). Equivalent Fos induction was seen regardless of the behavioral circumstance in which CNO was administered, with similar homecage mCherry-relative increases in hM3Dq homecage-tested rats (Sidak post hoc, *p* = 0.001) and hM3Dq rats tested in a novel operant chamber with (*p* = 0.0009) or without (*p* = 0.0004) cues. hM3Dq rats tested in operant boxes with or without cues, or tested in homecage did not differ in Fos expression (Sidak post hoc: *p*s > 0.99). These results collectively show that 1) hM3Dq stimulation augments neural activity as expected and 2) hM3Dq stimulation enhanced neural activity similarly regardless of the behavioral context in which the stimulation occurred.

### Inhibiting VP^GABA^ neurons suppresses remifentanil reinstatement after punishment

In Group^Punish^ rats expressing hM4Di DREADDs, a lever (active, inactive) x treatment (vehicle, CNO) ANOVA revealed a significant main effect of lever across all conditions (*p* < 0.01). Active lever presses in Context A with cues were suppressed by CNO treatment in hM4Di rats (Fig 4A, treatment x lever interaction: F_(1, 24)_ = 6.53, *p* = 0.017; active lever Sidak post hoc: *p* = 0.0047), but this was not the case in Context A without cues (Fig 4D, treatment: F_(1, 10)_ = 0.43, *p* = 0.52; treatment x lever interaction: F_(1, 10)_ = 3.38, *p* = 0.096), showing that inhibiting VP^GABA^ neurons suppressed seeking in Context A only in the presence of discrete cues. CNO treatment in hM4Di rats trended towards reducing opioid seeking in Context B with cues (Fig 4G, treatment x lever interaction: F_(1, 24)_ = 3.98, *p* = 0.058), but no main effect of treatment, or treatment x lever interaction was detected in Context B without cues (Fig 4J, treatment: F_(1, 24)_ = 2.79, *p* = 0.11; treatment x lever: F_(1, 24)_ = 0.12, *p* = 0.73).

**Figure 4.**
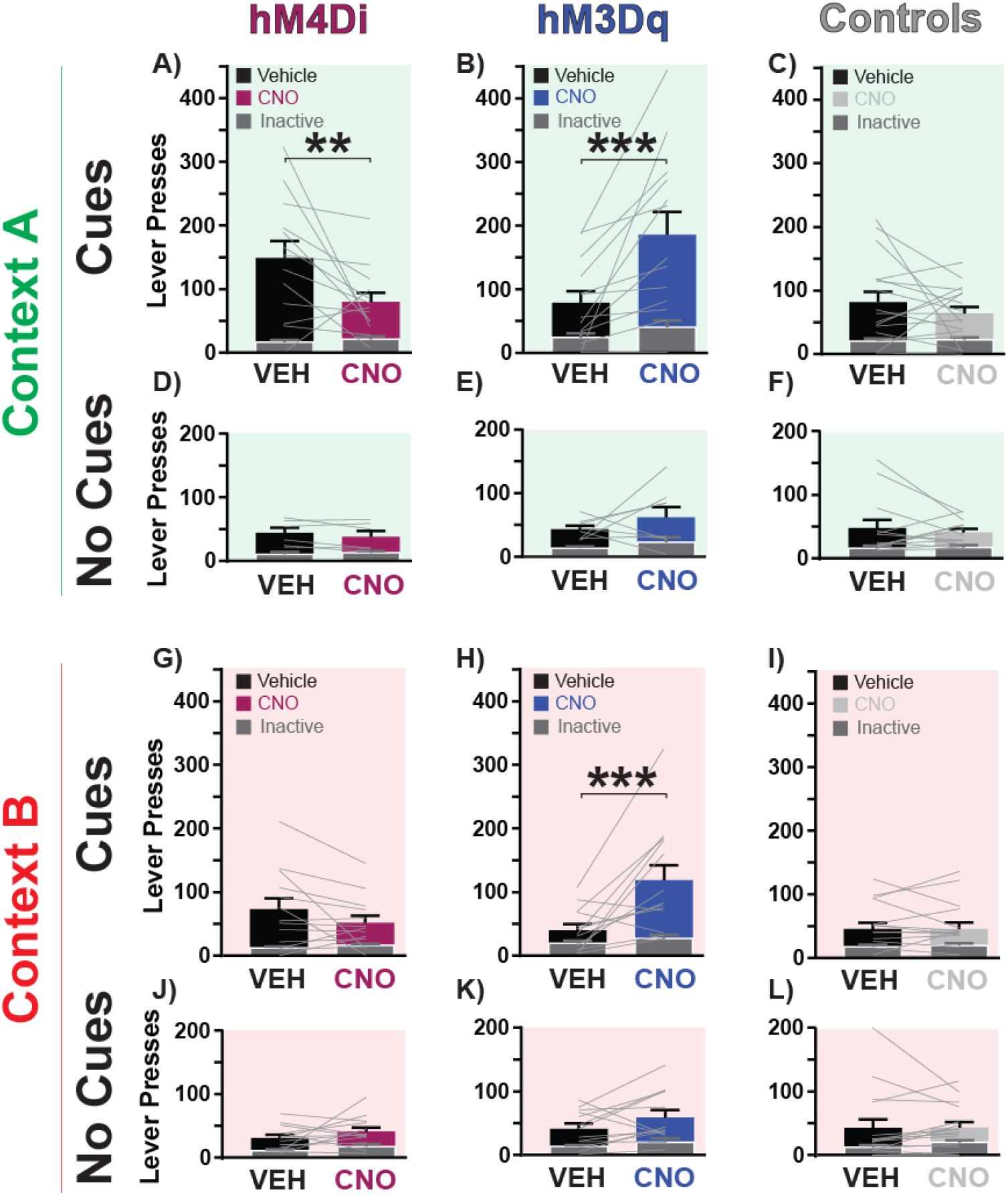
Following punishment, inhibiting or stimulating VP^GABA^ neurons bidirectionally controls remifentanil seeking. **A-C)** In Context A with cues, CNO treatment **A)** decreased opioid seeking in hM4Di rats (purple bar), **B)** increased seeking in hM3Dq rats (blue bar), and **C)** was without effect in control rats (light gray bar), relative to vehicle treatment (black bars). **D-F)** In Context A with no cues, CNO treatment was without effect on opioid seeking in D) hM4Di rats (purple bar), **E)** hM3Dq rats (blue bar), and **F)** control rats (light gray bar), relative to vehicle treatment day (black bars). **G-I)** In Context B with cues, CNO treatment **G)** was without effect on opioid seeking in hM4Di rats (purple bar), **H)** increased opioid seeking in hM3Dq rats (blue bar), and **F)** did not impact opioid seeking in control rats (light gray bar), relative to vehicle treatment day (black bars). **J-L)** In Context B with no cues, CNO treatment was without effect on opioid seeking in **J)** hM4Di rats (purple bar), **K)** hM3Dq rats (blue bar), and **L)** control rats (light gray bar), relative to vehicle treatment (black bars). Dark gray overlaid bars with white outline represent inactive lever presses, and light gray lines depict individual rats’ active lever pressing on each session. Repeated measures two-way ANOVA, Sidak post hoc: *p*\*\* < 0.01, *p*\*\*\* < 0.001. Data presented as mean ± SEM.

### Stimulating VP^GABA^ neurons augments remifentanil reinstatement after punishment

In Group^Punish^ rats expressing hM3Dq DREADDs, stimulating VP^GABA^ neurons strongly increased opioid seeking, and appeared to do so in a cue-dependent manner. Specifically, CNO treatment in hM3Dq rats augmented active lever pressing in both Context A with cues and Context B with cues (Fig 4B, Context A with cues treatment x lever interaction: F_(1, 24)_ = 8.78, *p* = 0.0068, active lever Sidak post hoc: *p* = 0.0001; Fig 4H, Context B with cues treatment x lever interaction: F_(1, 24)_ = 9.79, *p* = 0.0046, active lever Sidak post hoc: *p* = 0.0001). In contrast, CNO in hM3Dq rats failed to augment seeking in Context A in the absence of cues (Fig 4E, Context A with no cues treatment x lever interaction: F_(1, 14)_ = 0.22, *p* = 0.64; Context A with no cues treatment: F_(1, 14)_ = 2.08, *p* = 0.17). CNO in hM3Dq rats subtly increased pressing on both the active and inactive lever in Context B with no cues, as indicated by a main effect of treatment accompanied by a non-significant treatment x lever interaction (Fig 4K, Context B with no cues treatment: F_(1, 24)_ = 4.77, *p* = 0.039; Context B with no cues treatment x lever interaction: F_(1, 24)_ = 0.92, *p* = 0.35; active lever Sidak post hoc: *p* = 0.071; inactive lever: *p* = 0.63). For active lever timecourse details for each reinstatement/DREADD condition for Group^Punish^ rats, see Supplemental Figure 2.

### Stimulating VP^GABA^ neurons augments remifentanil seeking after extinction

In Group^Ext^ rats with hM3Dq DREADDs, CNO treatment augmented seeking in the presence of cues, irrespective of whether rats were in Context A or B (Fig 5A, treatment main effect Context A with cues: F_(1, 20)_ = 4.91, *p* = 0.038, active lever Sidak post hoc: *p* = 0.037; Fig 5E, treatment main effect Context B with cues: F_(1, 20)_ = 9.86, *p* = 0.0052, active lever Sidak post hoc: *p* = 0.033). In the absence of cues, CNO augmented opioid reinstatement only in Context A, but not in Context B (Fig 5C, treatment main effect Context A with no cues: F_(1, 20)_ = 8.84, *p* = 0.0075, active lever Sidak post hoc: *p* = 0.011 ; Fig 5G, treatment main effect Context B with no cues: F_(1, 20)_ = 2.10, *p* = 0.16; treatment x lever interaction: F_(1, 20)_ = 0.19, *p* = 0.67). Thus, stimulating VP GABA neurons in Group^Punish^ or Group^Ext^ rats augments seeking in either Context in the presence of cues, but only increases non-cued seeking in Context A in Group^Ext^ but not in Group^Punish^.

**Figure 5.**
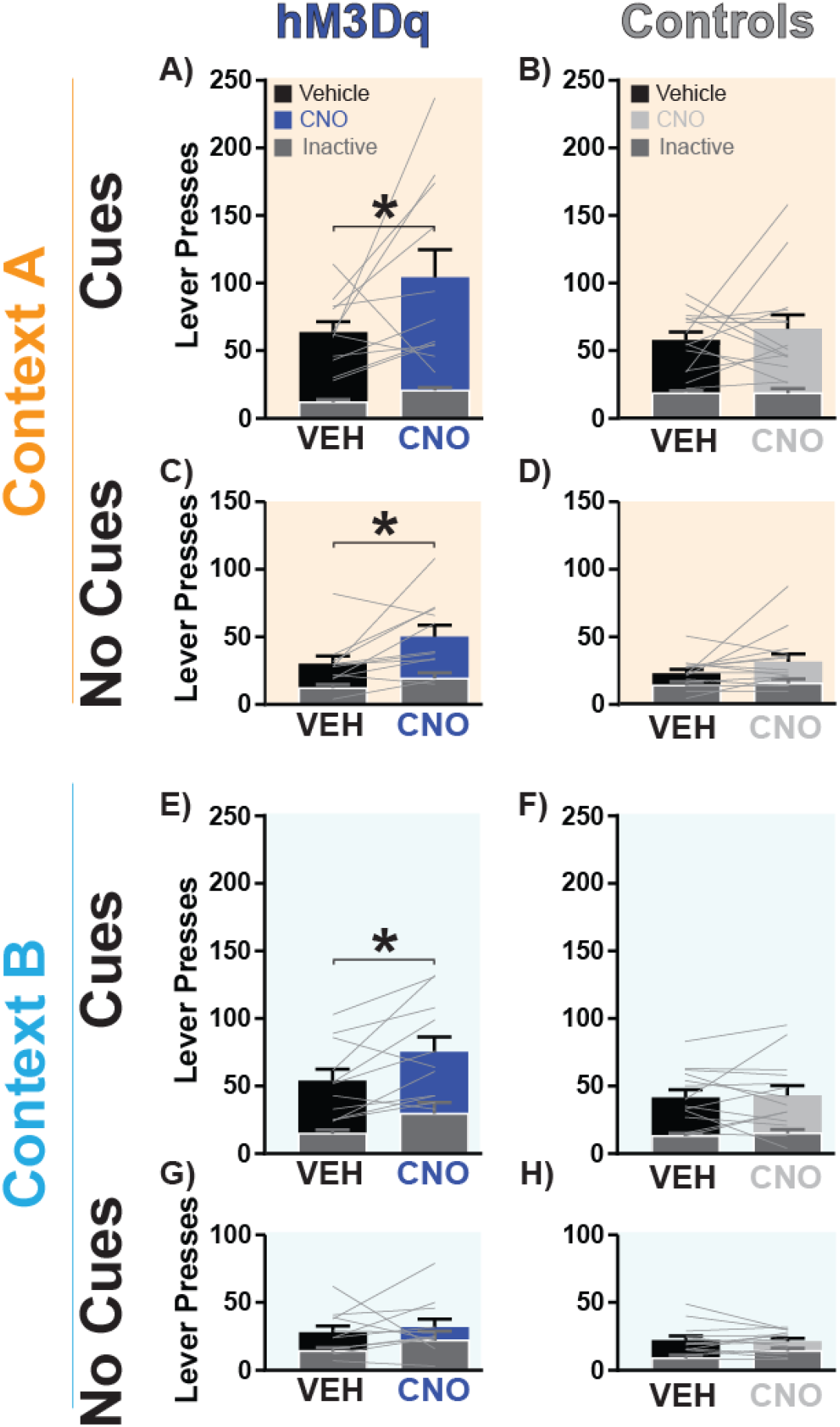
Following extinction, stimulating VP^GABA^ neurons augments reinstatement in a cue- and context-dependent manner. **A-B)** CNO treatment (blue bar) in hM3Dq rats augmented opioid seeking in Context A with cues relative to vehicle (black bar), but no effect was detected in controls (light gray bar versus black bar). **C-D)** CNO treatment (blue bar) in hM3Dq rats increased opioid seeking in Context A with no cues relative to vehicle, but no effect was seen in controls (light gray bar versus black bar). **E-F)** CNO (blue bar) in hM3Dq rats increased opioid seeking in Context B with cues, relative to vehicle treatment (black bar), but no effect was observed in controls (light gray bar versus black bar). **G-H)** CNO treatment in hM3Dq rats (blue bar) or controls (light gray bar) was without effect on opioid reinstatement in Context B with no cues, relative to vehicle treatment (black bars). Dark gray overlaid bars with white outline represent inactive lever presses, and light gray lines depict individual rats’ active lever pressing. *p*\* < 0.05. Data presented as mean ± SEM.

### Neither inhibiting nor stimulating VP^GABA^ neuron alters opioid self-administration

Finally, we asked whether VP^GABA^ neuron manipulations influence unpunished opioid self-administration in a subset of Group^Punish^ rats. Opioid self-administration was unaffected by CNO treatment in hM4Di rats (Fig 6A, treatment: F_(1, 10)_ = 2.14, *p* = 0.17; treatment x lever interaction: F_(1, 10)_ = 2.14, *p* = 0.17) or hM3Dq rats (Fig 6B, treatment: F_(1, 10)_ = 1.63, *p* = 0.23; treatment x lever interaction: F_(1, 10)_ = 1.63, *p* = 0.23).

**Figure 6.**
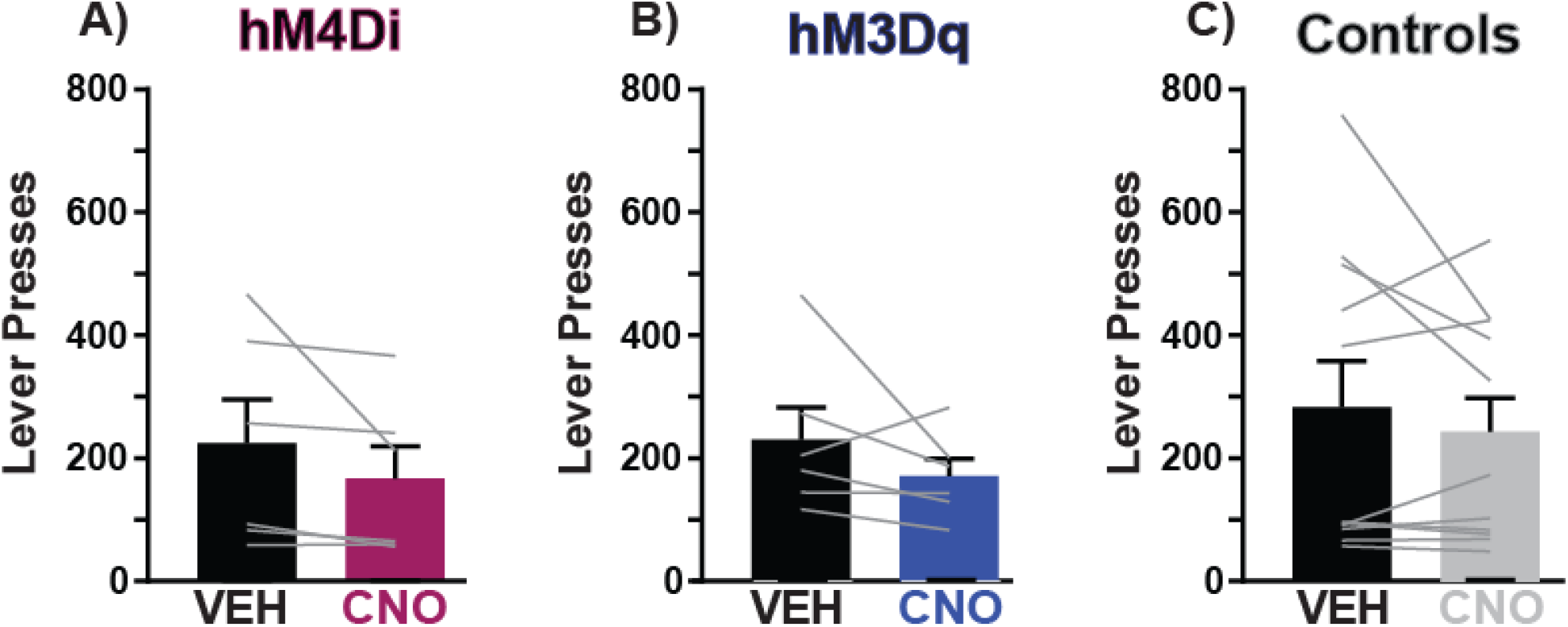
No impact of inhibiting or stimulating VP^GABA^ neurons on remifentanil self-administration. **A)** CNO treatment failed to alter unpunished (Context A) opioid self-administration in hM4Di rats (purple bar), **B)** hM3Dq rats (blue bar), or **C)** control rats (gray bar) relative to vehicle treatment (black bars). Light gray bars represent near-zero inactive lever presses (range 0 - 9 presses), and gray lines represent individual rats’ active lever pressing. Data presented as mean ± SEM.

### No effect of CNO on reinstatement or self-administration in control rats

In Group^Punish^ control rats, CNO did not influence opioid reinstatement in Context A with cues (Fig 4C) or without them (Fig 4F), or in Context B with (Fig 4I) or without cues (Fig 4L, treatment: *F*s < 0.86, *p*s > 0.35; treatment x lever interaction: *F*s < 1.28, *p*s > 0.26). Similarly, in Group^Ext^ control rats, CNO did not impact reinstatement after extinction in Context A with cues (Fig 5B) or with no cues (Fig 5D), or Context B with (Fig 5F) or with no cues (Fig 5H, treatment: *F*s < 3.56, *p*s > 0.06; treatment x lever interaction: *F*s < 2.61, *p*s > 0.11). CNO was also without effect on remifentanil self-administration in control rats (Fig 6C, treatment: F_(1, 20)_ = 0.98, *p* = 0.34; treatment x lever interaction: F_(1, 20)_ = 1.07, *p* = 0.31). Specificity of CNO effects was confirmed with two-way ANOVAs examining genotype (Group^Punish^: hM3Dq, hM4Di, controls) x treatment (vehicle, CNO) effects on active lever pressing in CNO-impacted reinstatement conditions. Significant interactions demonstrated CNO effects only in DREADD rats (Context A with cues: F_(2, 40)_ = 12.63, *p* < 0.0001, Context B with cues: F_(2, 39)_ = 13.34, *p* < 0.0001).

## Discussion

Using chemogenetic inhibition/stimulation and Fos expression analyses, we found that VP^GABA^ neurons play a key role in opioid relapse-like behavior. Following remifentanil self-administration and subsequent abstinence from drug taking, chemogenetically inhibiting VP^GABA^ neurons suppressed, and stimulation enhanced opioid reinstatement, especially when driven by discrete, response-contingent drug cues. VP^GABA^’s role was apparent across multiple reinstatement models, and it was specific to reinstatement, in that the same chemogenetic manipulations did not affect remifentanil self-administration. We also validated hM3Dq DREADDs as being capable of Fos-activating GABA neurons in GAD1:Cre transgenic rats, and determined that VP^GABA^ neurons were equivalently stimulated by DREADDs in the presence or absence of drug-associated cues, though the presence of cues during such stimulation was generally required for increased drug-seeking behavior. Finally, we found that endogenous neural activity (Fos) in rostral, but not caudal, VP correlated with reinstatement behavior. These experiments thus show a specific role for VP^GABA^ neurons in opioid relapse-like behaviors, regardless of the preclinical model employed—potentially positioning VP as a future target for intervention in this chronic, relapsing disorder.

In hopes of better modeling the circumstances of drug addiction, preclinical models have emerged in which drug taking is coupled with adverse consequences which cause rats to decide to quit taking drugs [57–60]—similar to the self-imposed abstinence present in most humans attempting to quit using drugs. We and others have suggested that through such efforts to better model human addiction and relapse-like behaviors in rats we may gain new insights into the neural circuit dynamics most likely engaged in addicted humans. Here, we build on prior work to establish a model of remifentanil cue- and context-induced relapse after punishment-induced abstinence, adapting those previously used with other drugs of abuse [51,61–64]. Using the short-acting but strongly reinforcing opioid drug remifentanil, we built on the work of Panlilio and colleagues, who previously established that footshock punishment suppresses remifentanil self-administration, and that remifentanil seeking can be subsequently reinstated [6,17]. Here, we expand on these models by incorporating an explicit contextual element to the reinstatement tests (with or without discrete drug-paired cues). We note that unlike extinction- or forced abstinence-based reinstatement models, voluntary abstinence models may mimic the conflicted motivational processes that often arise in addicted people attempting to control their drug use due to mounting life consequence. We hope that developing this approach in rats this could ultimately lead to deeper understanding of neural circuits that are engaged when humans decide whether or not to pursue drugs.

Discrete cues occurring in conjunction with drug use (e.g., paraphernalia), and diffuse contextual elements (eg, location of prior drug use) serve as powerful triggers that can ultimately lead an abstinent person to relapse. The ability of discrete cues and contexts to elicit drug seeking appear to depend on overlapping yet distinct neural circuits [59,65,66], some of which involve VP or its close neural connections [33,47,67–72]. Therefore, we examined VP^GABA^ involvement in reinstatement elicited by both discrete cues and also contexts in our behavioral relapse models. After punishment-induced abstinence, inhibiting VP^GABA^ neurons only reduced remifentanil seeking in the “safe” Context A in the presence of cues—the condition in which reinstatement was highest. In contrast, we found that inhibiting VP^GABA^ neurons did not affect seeking in the punishment-associated Context B in the presence or absence of cues, potentially in part due to a floor effect resulting from low responding in this “dangerous” context. These results are reminiscent of our prior report with cocaine showing that chemogenetic inhibition of VP neurons suppressed cue-induced drug seeking in a safe Context A, but not in a dangerous Context B, using an analogous voluntary abstinence-based reinstatement model [51]. Here, we also examined effects of stimulating VP^GABA^ neurons on post-punishment opioid seeking, which we found to robustly augment cue-induced remifentanil seeking in both Context A and Context B. In the absence of cues, however, stimulating VP^GABA^ neurons exhibited no effect in either Context A or B. It appears, then, that response-contingent cues are required to reveal the motivation-enhancing effects observed with hM3Dq stimulation, at least after punishment-induced abstinence. Overall, these data suggest cue- and context-dependent roles for VP GABA in reinstatement following voluntary abstinence.

Though we found cue- and context-dependent effects of manipulating VP^GABA^ neurons on opioid seeking following punishment-induced abstinence, the way in which abstinence is achieved in preclinical models determines the neural circuits recruited during reinstatement [7,58,59,73]. Therefore, we asked whether stimulating VP^GABA^ neurons would have similar effects on cue or context-induced reinstatement using an analogous extinction-based abstinence reinstatement model. We found that VP^GABA^ neuron stimulation in extinguished rats similarly augmented cue-induced remifentanil seeking in both Context A and B. However, unlike in punishment-trained rats, VP^GABA^ neuron stimulation in extinguished rats augmented seeking in Context A in the absence of cues, not just in their presence. This could suggest that VP^GABA^ roles in context-induced reinstatement may differ based on the affective associations imbued in these contexts (i.e. fear of shock versus extinction-related disengagement), or contrast effects between the always safe Context A with the extinction- or punishment-paired Context B. Alternatively, differences between the models in the number of training days required for extinction-versus punishment-induced abstinence (Fig S1), or other methodological differences between the procedures could have contributed to this distinction. Regardless, these findings of a relatively pervasive, necessary and sufficient role for VP^GABA^ neurons in cue-induced relapse contrast with other VP-connected limbic nodes like the basolateral amygdala or dorsal striatum, since inactivating these nodes differentially affects reinstatement behavior depending on the way in which abstinence was achieved [7,73]. Overall, these results suggest that VP^GABA^ neurons are involved in cue-induced drug seeking across rat relapse models, suggesting they might also play a pervasive role in opioid relapse in humans.

Prior work from us and others suggests nuanced roles for VP and its neuronal subpopulations in drug seeking, as well as motivated behavior more generally [32,35,50,74–76]. In particular, recent reports support a role for phenotypically-defined VP cellular subpopulations in relapse to drug seeking across drugs of abuse and relapse models [29,33,51,67,77–80]. For example, VP dopamine D3-receptor expressing populations and their outputs to lateral habenula are critical for cue-induced cocaine seeking [77]. VP^GABA^ and parvalbumin-expressing VP neurons are also recruited by alcohol-associated contextual cues, and chemogenetic inhibition of these subpopulations suppressed context-induced relapse to alcohol seeking [47]. Stimulation of VP^GABA^ neurons also enhanced extinction responding after cocaine self-administration, and stimulation of a subset of enkephalin-expressing VP^GABA^ neurons enhanced cue-induced cocaine reinstatement, whereas stimulating VP glutamate neurons instead suppressed cocaine reinstatement [29]. These findings are in accordance with the idea that VP glutamate neurons constrain reward seeking, and have opposite motivational roles to VP^GABA^ neurons, which are instead involved in appetitive processes [27,28,30,50]. Our results further demonstrate the critical role of VP^GABA^ neurons in opioid seeking, especially when triggered by drug-paired cues.

Consistent with prior reports examining cocaine seeking, we identified that rostral, but not caudal, VP neural activity (Fos) was positively correlated with cue-induced drug seeking [33], and that VP neural activity was elevated following reinstatement testing in both punishment-associated Context B or reward-associated Context A [51]. Our Fos results demonstrate that rostral, not caudal VP is activated when rats undergo cue-induced reinstatement of remifentanil seeking, and that rostral VP Fos scales with the intensity of drug seeking across individual animals. Several groups have also shown a functional gradient along VP’s rostrocaudal axis [36,38–42]. For example, caudal VP contains a ‘hedonic hotspot’ in which locally applied orexinA or μ opioid receptor agonists enhance hedonic orofacial ‘liking’ reactions to sweet liquid rewards [34,43]. Though our Fos analysis was not restricted to VP^GABA^ cells, the majority of Fos-positive cells were likely GABAergic, since VP consists of mostly GABAergic neurons across its rostrocaudal axis [27,32,81]. We also note that our DREADD manipulations were targeted in central VP, and therefore spanned both rostral and caudal VP zones. Future work should further dissect anatomical, cellular, and molecular profiles spanning rostrocaudal VP zones to determine the specific roles of neuronal populations within these subregions responsible for this apparent anterior-posterior functional gradient.

It was clear that Gq-DREADDs robustly stimulate Fos in VP^GABA^ neurons, as seen in other neural populations [33,53,82–85], and here we further asked whether the behavioral situation impacts hM3Dq-induced Fos, as it does for hM3Dq-induced behavior. Specifically, we reasoned that since hM3Dq-stimulated reinstatement was most robust in the presence of response-contingent cues, the presentation of these cues might further augment hM3Dq-simulated Fos levels, relative to rats exposed to no cues, or tested outside a drug-seeking context (homecage). However, the presence or absence of discrete, passively administered cues made no difference—Fos levels in hM3Dq-mCherry VP neurons following CNO administration were similar regardless of whether testing occurred in the presence of drug cues. This indicates that hM3Dq DREADD stimulation enhances activity of VP^GABA^ neurons regardless of behavioral situation, though the same stimulation only caused consistent increases in drug seeking the presence of cues. This likely implies that VP activation either results in drug seeking or does not depending on the cue-elicited activity state of the wider motivation circuits within which VP interacts. Perhaps this is not surprising—enhanced activity of the VP^GABA^ projection neurons cannot inhibit cue-evoked activity in target regions (e.g. VTA) if cues are not present to cause activity capable of being inhibited. Conceptually, this could be a useful feature of manipulating such inhibitory circuits to treat psychiatric disorders, as the behavioral/cognitive effects of activating GABAergic projections may only become apparent in the presence of symptom-related circumstances related to abnormal neuronal hyperactivity in downstream regions.

Our results establish that VP^GABA^ neurons critically regulate reinstatement across preclinical opioid relapse models, adding to the growing evidence for a key role of VP within motivation circuits [19,20,32,86–92]. Though such evidence implies that targeting VP circuits might be a useful strategy for helping humans struggling to control their drug use, many questions remain. How does molecular heterogeneity within neurotransmitter-defined VP circuits influence motivated behavior? Is it possible to modulate the activity of VP to influence maladaptive drug seeking without impairing healthy desires? How is VP’s efferent and afferent connectivity involved in different types of motivated behavior? Further investigating these questions may yield fruitful insights not only about VP’s role in addiction, but also fundamental ways in which motivational systems interact with cognitive and memory systems more generally.

---

**Supplemental Figure 1.**
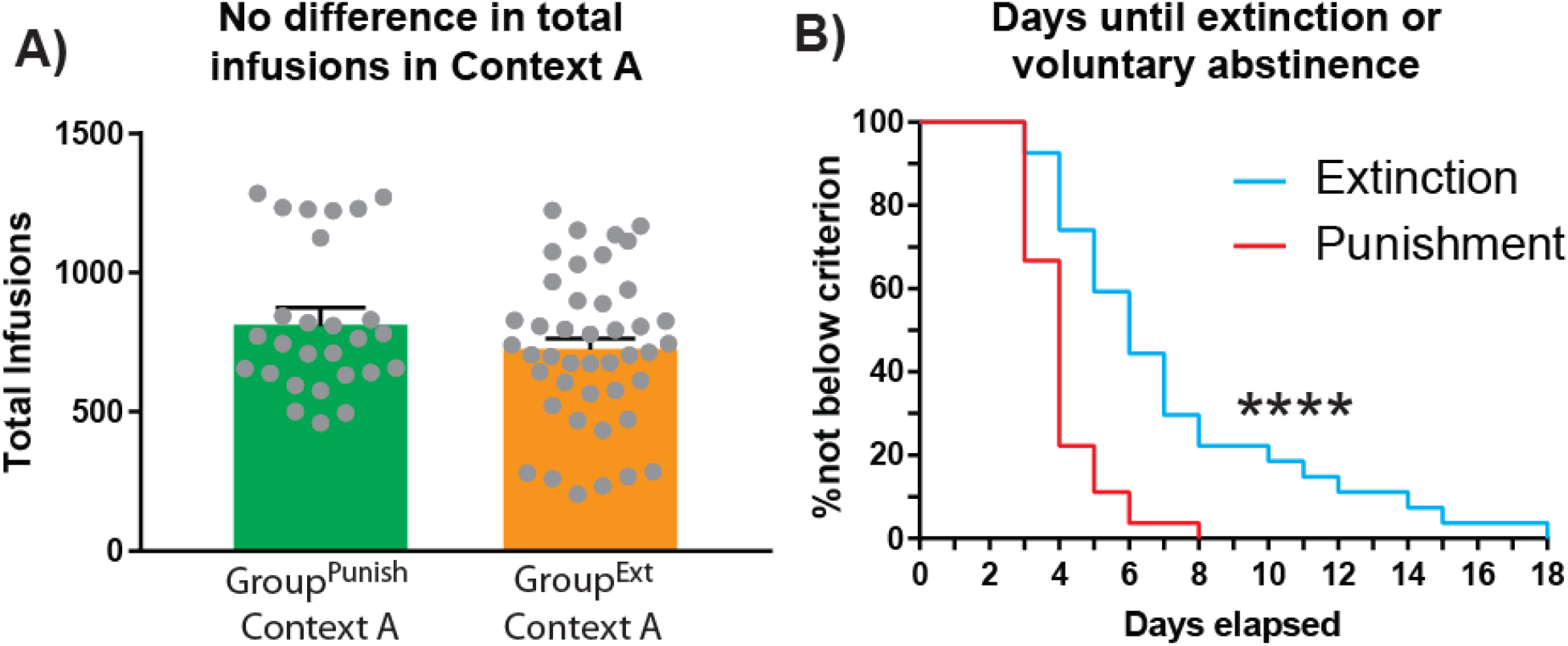
Comparison of Context A and Context B behavior across punishment and extinction models. **A)** No difference in infusions earned during 14-day Context A training for rats that subsequently underwent punishment training (green bar) or extinction (orange bar). Individual data points presented as gray dots. **B)** Survival analysis shows that Group^Ext^ rats in who underwent extinction training in Context B (blue line) spent more days above criterion (< 25 active lever presses on 2 consecutive days), relative to Group^Punish^ rats that underwent punishment training in Context B (red line). Log-rank (Mantel-Cox) test, *p*\*\*\*\* < 0.0001.

**Supplemental Figure 2.**
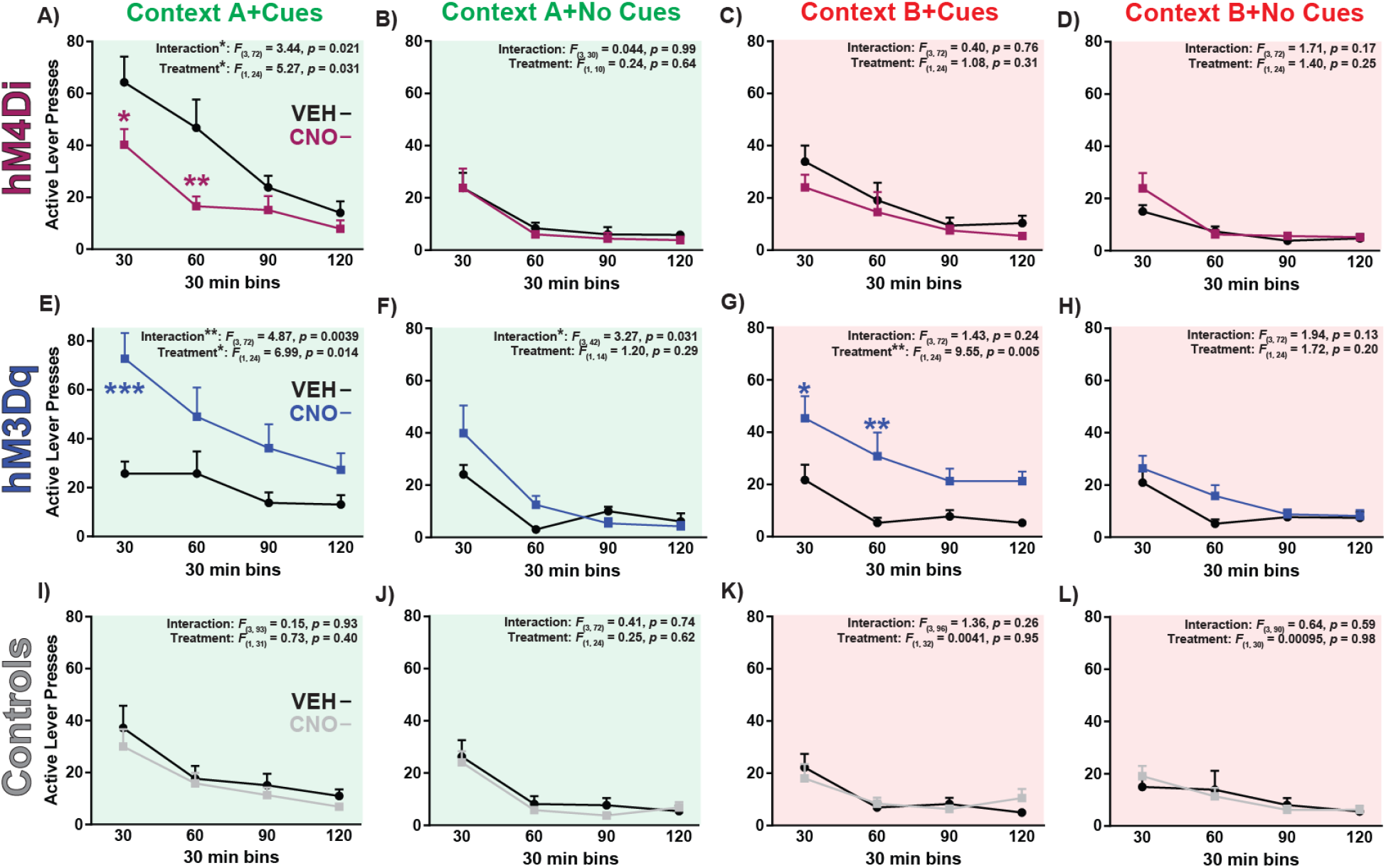
Group^Punish^ hM4Di, hM3Dq, and control rats’ reinstatement timecourse following punishment training. **A-D)** A two-way ANOVA with treatment and time block as factors revealed a significant main effect of CNO treatment and treatment x time block interaction in hM4Di rats in Context A with cues. Colored asterisks show significant Sidak post hoc tests between CNO and vehicle (*p*\* < 0.05, *p*\*\* < 0.01, *p*\*\*\* < 0.001). Two-way ANOVA revealed no significant main effect or treatment x time block interaction in hM4Di rats in **B)** Context A with no cues, **C)** Context B with cues, or **D)** Context B with no cues. **E)** Two-way ANOVA revealed a significant main effect of CNO treatment and treatment x time block interaction in hM3Dq rats in Context A with cues. **F)** Two-way ANOVA revealed a significant treatment x time block interaction in hM3Dq rats in Context A with no cues. **G)** Two-way ANOVA revealed a significant main effect of treatment, with no treatment x time block interaction in hM3Dq rats in Context B with cues. **H)** No main effect of treatment or significant treatment x time block interaction was detected for hM3Dq rats in Context B with no cues. **I-L)** Two-way ANOVAs revealed no main effect of treatment or significant treatment x time block interaction for **I)** Context A with cues, **J)** Context A with no cues, **K)** Context B with cues, or **L)** Context B with no cues. All conditions exhibited a significant main effect of time block (*F*s > 7.35, *p*s < 0.0003). Purple, gray, and blue lines represent CNO treatment and black lines represent vehicle treatment. Data presented as mean ± SEM.

